# Effects of ocean acidification on red king crab larval survival and development

**DOI:** 10.1101/2023.10.02.560246

**Authors:** W. Christopher Long, Allie Conrad, Jennifer Gardner, Robert J. Foy

## Abstract

Ocean acidification, a decrease in oceanic pH resulting from the uptake of anthropogenic CO2, can be a significant stressor for marine organisms. In this study, we reared red king crab larvae from hatching to the first crab stage in four different pH treatments: current surface ambient, diel fluctuation to mimic larval migration between the surface and mixed layer under current ambient conditions, pH 7.8, and pH 7.5. Larvae were monitored throughout development and the average length of each stage was determined. At each of the zoeal stages, the glaucothoe stage, and the first crab stage, we measured survival, morphometry, dry mass, and carbon, nitrogen, calcium, and magnesium content. Red king crab larvae were highly resilient to ocean acidification. There were no differences among treatments in survival or in average stage length. Although there were clear ontogenetic trends in size, weight, and elemental composition, most of these did not vary with pH treatment. Zoeal morphology did not vary among treatments, although glaucothoe and C1 crabs were slightly smaller in pH 7.8 than in the ambient treatment. Ambient larvae also had a slightly higher mass than pH 7.8 larvae but not pH 7.5. Ambient larvae had higher magnesium contents than pH 7.8 and pH 7.5, but calcium levels were the same. Ambient larvae also had slightly lower carbon and nitrogen content than pH 7.8 and pH 7.5 larvae but only in the 4th zoeal stage. Overall this study suggests that red king crab larvae are well adapted to a wide range of pH conditions and are unlikely to be significantly affected by ocean acidification levels projected for the next two centuries.

## Introduction

Increasing CO_2_ in the atmosphere, caused by human emissions, is causing a commensurate increase in the CO2 in the oceans which directly affects the carbonate chemistry and results, among other things, in a decrease in the pH of the water known as ocean acidification. Average global pH had decreased about 0.1 units since the start of the industrial revolution and a further 0.3-0.5 unit change is expected by the end the century. Many, but not all, marine species are negatively affected by such a reduction in pH and ocean acidification is therefore likely to cause shifts in marine ecosystems.

Decapod crustaceans, similar to other taxa, have a range of responses to reduced pH both within (Walther et al., 2010; Long et al., 2013b; Long et al., 2016) and among (Bednaršek et al., 2021) species. Decreased water pH causes acute acidosis of crustacean hemolymph which many, but not all species, are able to respond to by buffering their hemolymph via active ion transport (Pane and Barry, 2007; Small et al., 2010) or by reducing their metabolic rates and thus the metabolic production of CO_2_ (Pane and Barry, 2007; Small et al., 2020). These responses can bring the hemolymph back into acid-base homeostasis (Meseck et al., 2016); however, the energetic costs of these processes can shift energy investment away from other processes such as growth and reproduction (Small et al., 2016; Swiney et al., 2016).

Red king crab in the Bering Sea are an important, commercially harvested species. Red king crab have a complex life history. Female red king crab brood ∼50,000-450,000 embryos (Swiney et al., 2012; Swiney and Long, 2015) for about a year and larvae hatch in the spring. The larvae are planktonic and pass through four feeding zoeal stages before molting to the settling, non-feeding glaucothoe stage (Nakanishi, 1987; Kovatcheva et al., 2006). Glaucothoe are semi-demersal and actively search for complex habitat (Stevens and Kittaka, 1998; Stevens, 2003); both glaucothoe (Stevens and Swiney, 2005) and early benthic stage juveniles (Long et al., 2012) are dependant on complex habitat because they avoid predation (Long et al., 2015; Lyons et al., 2016) both from con-specifics (Long and Whitefleet-Smith, 2013) and other species through crypsis (Pirtle et al., 2012; Daly et al., 2013). At about 2 years they become too large to effectively hide from predation and start to exhibit podding behavior, analogous to schooling in fish, to avoid predation (Powell and Nickerson, 1965; Dew, 1990). As they grow they move into deeper, more unstructured habitats where they grow to maturity (Dew, 1990).

Red king crab are sensitive to ocean acidification. Exposure during the embryo stage alters development and hatch timing (Long et al., 2013a). First stage larvae have decreased starvation-survival time, and carryover effects from embryo exposure are negative (Long et al., 2013a). Juveniles suffer decreased growth and increased mortality at a pH of 7.8 (Long et al., 2013b) and increased temperature has a synergist negative effect (Swiney et al., 2017). Right after exposure to low pH, juvenile respiration increases; however, feeding levels do not increase (Long et al., 2019). This suggests a high energetic cost of responding to low pH without an equivalent increase in energy intake. Adults and first stage larvae calcify faster in response to low pH (Long et al., 2013a) and juveniles maintain calcification levels (Long et al., 2013b); however, despite this, changes to the cuticle lead to thinning of the carapace and decreased hardness in the chelas in juveniles at low pH (Coffey et al., 2017). These effects are predicted to have a negative effect on the red king crab population and fisheries in Alaska in the near future (Punt et al., 2014; Punt et al., 2022) Nothing is known about the response of larvae past the first stage to low pH, so, in this study, we reared red king crab larvae from hatching to the first crab stage in four different pH treatments to fill this critical gap.

## Methods

### Experimental design and larval rearing

Ovigerous red king crab were collected from Bristol Bay and transported to the Kodiak laboratory. Crabs were held in 2000 L tanks with flowing ambient seawater and fed to excess twice per week. Embryos were monitored and crabs were moved to 48 L individual holding tanks when embryos approached hatching. Hatching was monitored by capturing larvae in nets until the majority of crabs were hatching. Larvae were collected overnight and experimental larval tanks stocked in a random order (see below for treatments). Larval tanks were stocked over a three-day period; nine tanks in each of the first two days and two tanks in the final day. Each individual tank was fully stocked in a single day. Larvae were collected from the same 21 females each of the three days; however, on the third day one of the females had stopped hatching. On each day of stocking, the larvae from all the females was pooled and the total number of larvae estimated by gently stirring the water the larvae were in until the larvae were distributed evenly. Subsamples of known volume were then removed and the number of larvae in each was counted and the average concentration of larvae was determined and used to estimate the total number of larvae collected. To stock each larval tank, the volume of water needed to stock each tank was calculated using the concentration of larvae, removed, placed into a separate container, and brought up to a known volume. Then the total number of larvae was estimated volumetrically, as above, to determine starting number of larvae in each tank and then the larvae were transferred to each tank.

Experimental larval tanks were circular with a conical bottom and had a volume of 180 L. Sand-filtered incoming seawater at ambient temperature and salinity was passed through a 5 µm mesh bag filter and UV sterilized. Each tank received flow of 2 L per minute. Air stones at the base of the standpipe were used to circulate water within each tank. Each of 20 tanks was randomly assigned to one of four pH treatments (five per treatment). Three had a constant pH: Ambient (ambient surface pH ∼8.05), pH 7.8 (pH expected in global surface waters in ∼2100), and pH 7.5 (∼2200). These pH treatments were used also, in part, to allow direct comparisons between this experiment and previous experiments on king crab (Long et al., 2013b; Long et al., 2017; Long et al., 2019). Because there is evidence that zoeal king crab may undergo a diel vertical migration (Haynes, 1983; McMurray et al., 1986; Wainwright et al., 1991) we included a treatment, hereafter referred to as Diel where pH was kept at surface ambient during the night and reduced by 0.1 pH units during the day to simulate movement to the mixed layer during the day under current conditions (Mathis et al., 2011) during the zoeal stages of larvae development.

pH was adjusted by bubbling pure CO_2_, the flow of which was controlled by Honeywell controllers and Durafet III pH probe, into each experimental tank. pH and temperature were measured daily in all tanks and twice daily in diel tanks using a Durafet III pH probe calibrated daily with TRIS buffer (Millero, 1986). Once a week, a water sample was taken from each tank, poisoned with mercuric chloride, and analyzed for total dissolved inorganic carbon (DIC) and total alkalinity (TA). DIC and TA were determined using a VINDTA 3C (Marianda, Kiel, Germany) coupled with a 5012 Coulometer (UIC Inc.) according to the procedure in (DOE, 1994) using Certified Reference Material from the Dickson Laboratory (Scripps Institute, San Diego, CA, USA; (Dickson et al., 2007)). The other components of the carbonate system were calculated in R (V3.6.1, Vienna, Austria) using the seacarb package (Lavigne and Gattuse, 2012). Water temperatures increased throughout larval rearing from 4.8 °C in early April when the experiment started to 9.4 °C in early July when it finished. The pHs averaged within 0.01 units of the nominal values and the pH 7.8 treatment was undersaturated with respect to aragonite and the pH 7.5 undersaturated with respect to calcite (Table 1)

**Table 1:** Physical and chemical properties of seawater in red king crab larval tanks. Temperature and pH were measured daily, dissolved inorganic carbon (DIC) and alkalinity were measured weekly, and all other parameter were calculated. Values are mean ± one standard deviation.

|  | Ambient | Diel AM | Diel PM | pH 7.8 | pH 7.5 |
| --- | --- | --- | --- | --- | --- |
| **Temperature** | 7.24 ± 1.40 | 7.22 ± 1.40 | | 7.25 ± 1.40 | 7.23 ± 1.44 |
| **Salinity** | 31.267 ± 0.142 | 31.268 ± 0.133 | 31.317 ± 0.124 | 31.277 ± 0.149 | 31.288 ± 0.163 |
| **pH<sub>T</sub>** | 8.05 ± 0.03 | 7.98 ± 0.10 | 8.06 ± 0.03 | 7.79 ± 0.05 | 7.50 ± 0.06 |
| **pCO<sub>2</sub> μatm** | 370.74 ± 26.92 | 467.73 ± 50.84 | 383.22 ± 23.62 | 703.89 ± 90.45 | 1414.71 ± 287.82 |
| **HCO<sub>3</sub><sup>-</sup> mmol/kg** | 1.89 ± 0.08 | 1.96 ± 0.07 | 1.91 ± 0.08 | 1.96 ± 0.05 | 2.00 ± 0.04 |
| **CO<sub>3</sub><sup>-2</sup> mmol/kg** | 0.11 ± 0.01 | 0.09 ± 0.01 | 0.11 ± 0.01 | 0.06 ± 0.01 | 0.03 ± 0.01 |
| **DIC mmol/kg** | 2.01 ± 0.08 | 2.04 ± 0.08 | 2.04 ± 0.09 | 2.06 ± 0.05 | 2.10 ± 0.05 |
| **Alkalinity mmol/kg** | 2.09 ± 0.08 | 2.08 ± 0.07 | 2.07 ± 0.08 | 2.10 ± 0.05 | 2.11 ± 0.04 |
| **$\Omega_{\text{Aragonite}}$** | 1.66 ± 0.09 | 1.41 ± 0.15 | 1.62 ± 0.08 | 0.96 ± 0.13 | 0.52 ± 0.19 |
| **$\Omega_{\text{Calcite}}$** | 2.65 ± 0.15 | 2.24 ± 0.24 | 2.58 ± 0.13 | 1.53 ± 0.21 | 0.83 ± 0.31 |

Larval rearing followed standard red king crab culturing techniques (Swingle et al., 2013; Long, 2016). In brief, zoea were fed daily to excess on a diet of *Artemia* nauplii enriched with DC DHA Selco (INVE Aquaculture, Salt Lake City). Uneaten artemia were flushed out of the tanks each day prior to feeding. To reduce bacterial growth, 1.9g of EDTA dissolved in seawater was added to each tank every day. When all of the larvae in a tank had molted to the glaucothoe stage structure, in the form of artificial seaweed, was added to each tank to give the glaucothoe a substrate to settle onto. The experiment was ended after all glaucothoe had molted to the first crab (C1) stage.

### Data collection

Each day, five larvae were removed from each tank and the developmental stage was determined. Once each molt stage, five days after the first day that 100% of the sampled larvae molted to that stage, 50 larvae were counted out, dried to a constant mass at 60 C and the average dry mass was calculated. At the same time, ∼250 larvae were removed from the tank and dried. They were homogenized. A subsample was analyzed for carbon and nitrogen (CN) content using the Dumas combustion method and an automated organic elemental analyzer (Gnaiger and Bitterlich, 1984). A second subsample was analyzed for calcium and magnesium content via inductively coupled plasma-atomic emission spectrometry. Four to seven days after each molt the tank was drained and the larvae carefully collected. The number of surviving larvae was estimated volumetrically as above and the percent survival, both from the previous stage and cumulatively was calculated. For one tank during the count at the Z2 stage most of the larvae were accidentally lost; survival data from this tank was not included in any analysis. At the same time, to measure morphometrics on the larvae, three larvae were removed and photographed under a microscope. On zoeal stages, carapace height, carapace length, total length, rostrum length, and telson spine length were measured using image analysis software as per Long et al. (2013a). On glaucothoe and C1 stages, carapace width, carapace length, rostrum base width, orbital spine width, and the first spine length were measured as per Long et al. (2013b).

### Analysis

The staging data was fit to a multi-stage transition model as per Long (2016). In brief each molt from one stage to the next is modeled as: 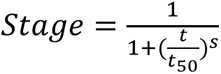, where t is the time in degree days (to account for the effect of temperature on the development of ectotherms), *t_50_* is the time at which 50% of the individuals have molting to the next stage and *s* is proportional to the abruptness of the transition. The data from each tank was fit separately resulting in a *t_50_* and *s* estimate for each stage in each tank. We calculated the stage duration for each stage as the time between *t_50_* estimates for each tank. Further, we calculated the slope at each transition as 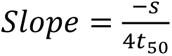 as a measure of the rapidity of the transition between the stages (Long, 2016). Both the stage duration and the *Slope* estimates were analyzed with an ANOVA with treatment fully crossed with larval stage and tank (nested within treatment) as factors. In addition, to ensure that potential additive effects across stages were not missed, the cumulative degree days to each stage, the *t_50_*, was analyzed with one-way ANOVAs with treatment as the factor.

Percent survival between subsequent stages was analyzed with an ANOVA with treatment fully crossed with larval stage and tank (nested within treatment) as factors. Cumulative survival to each stage was analyzed with a one-way ANOVA with treatment as the factor. Dry mass and C, N, Ca, and Mg contents were analyzed with an ANOVA with treatment fully crossed with larval stage and tank (nested within treatment) as factors.

Morphometrics for the zoeal stages were analyzed separately from those of the G and C1 stages because ontogenetic differences in organismal shape meant that the measurements taken on the two groups were different and could not be compared. In both cases, the data was normalized (expressed in terms of its z-value) and was visualized with non-metric multidimensional scaling (nMDS) plots and analyzed with permutational multivariate analysis of variances (PERMANOVA) with treatment fully crossed with larval stage and tank (nested within treatment) as factors from a Euclidian distance resemblance matrix.

## Results

The average stage duration increased with larval stage (Table 2, Fig. 1) with all stages differing from all other stages except Z2 and Z3 were the same (Tukey’s test, p < 0.0005 in all cases), but did not differ among treatments Table 2) or their interaction (Table 2). The cumulative time to each stage did not differ among any treatments (p > 0.19 in all cases) except for the molt to glaucothoe (F_3,16_ = 3.861, p = 0.03) where larvae in the pH 7.8 treatment took 13.9 degree days (effect size 4.5%) longer than larvae in the Ambient treatment (Fig. 1).

**Figure 1:**
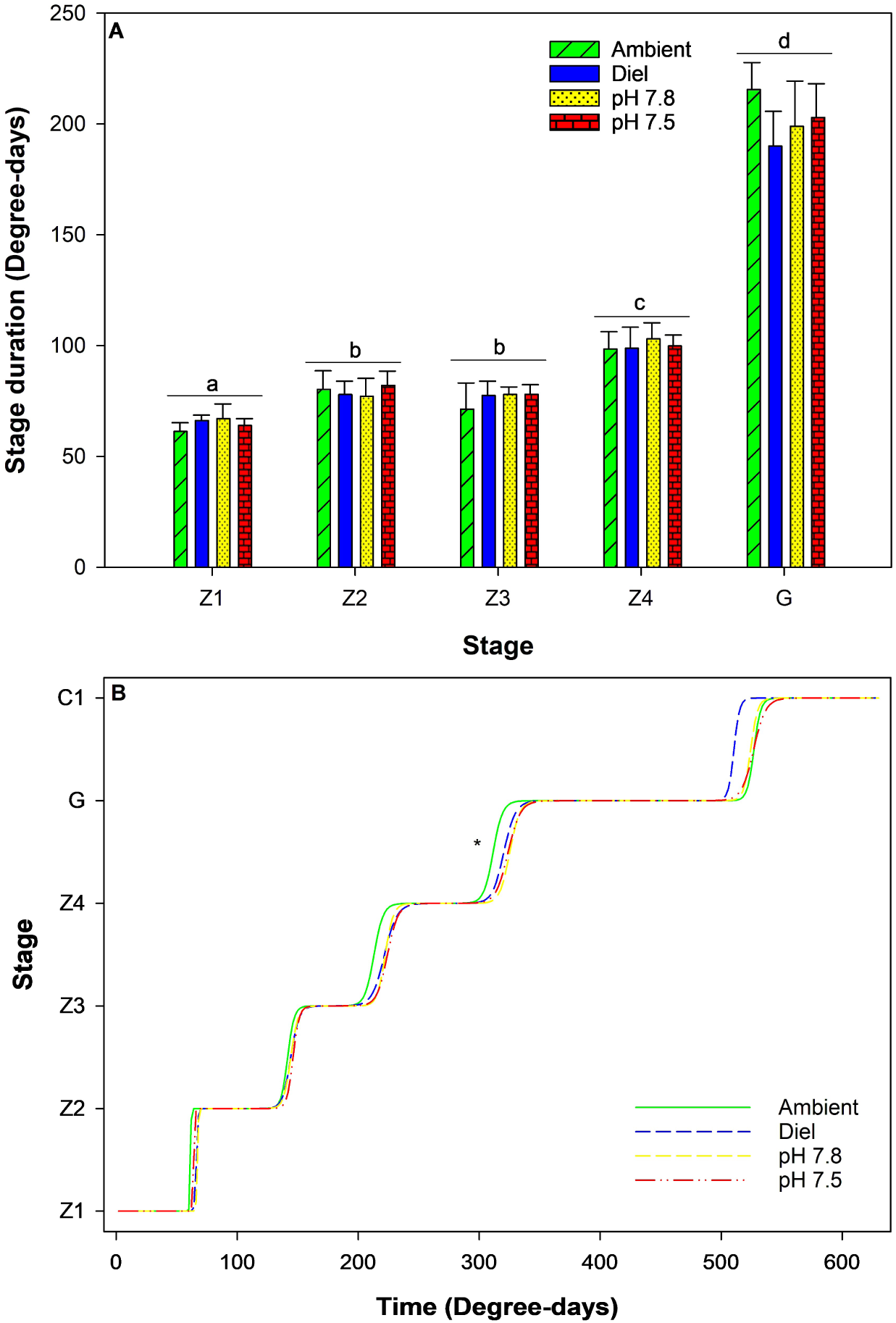
Red king crab larval development from hatching to the first crab stage in four different pH treatments. A) Development time for each stage. Bars with different letters above them are significantly different. Bars are means plus one standard deviation. B) Cumulative development of red king crab larvae. Lines represent best-fit multiple stage transitions models averaged within each pH treatment. Star indicates a significant difference among treatments in cumulative degree days at the molt to the glaucothoe stage.

**Table 2:** Summary ANOVA statistics for red king crab larval experiments. Treatment represents the experimental pH treatment and stage the developmental stage of the larvae. Response include the duration of each stage (stage duration) in degree days; the slope of the multiple-stage transitions model at the t_50_ for each stage (slope); percent survival; average larval mass; and the carbon, nitrogen, calcium and magnesium contents of the larvae (C, N, Ca, and Mg) as percent dry mass.

|  | Treatment |  | Stage |  | Treatment * Stage |  | Tank(Treatment) |  |
| --- | --- | --- | --- | --- | --- | --- | --- | --- |
|  | F | p | F | p | F | p | F | p |
| **Stage duration** | 0.645 | 0.589 | 652.557 | < 0.0005 | 1.644 | 0.102 | 0.592 | 0.878 |
| **Slope** | 0.33 | 0.806 | 12.48 | < 0.0005 | 0.30 | 0.988 | 1.07 | 0.405 |
| **Survival** | 0.14 | 0.935 | 27.88 | < 0.0005 | 1.93 | 0.049 | 0.78 | 0.699 |
| **Mass** | 2.98 | 0.042 | 186.24 | < 0.0005 | 1.74 | 0.093 | 1.85 | 0.058 |
| **C** | 3.39 | 0.026 | 236.38 | < 0.0005 | 2.06 | 0.041 | 1.14 | 0.354 |
| **N** | 2.91 | 0.045 | 368.37 | < 0.0005 | 2.39 | 0.018 | 1.37 | 0.204 |
| **Ca** | 1.19 | 0.326 | 114.48 | < 0.0005 | 0.76 | 0.689 | 1.67 | 0.095 |
| **Mg** | 5.19 | 0.004 | 30.82 | < 0.0005 | 1.31 | 0.25 | 1.11 | 0.375 |

Survival differed among the stages with the molt from the glaucothoe stage to the first crab stage having the lowest survival but did not differ among treatments (Table 2, Fig. 2). There was a significant interactive effect (Table 2); however post-hoc comparisons showed no significant differences among treatments within any stage (Tukey’s test, p > 0.099 in all cases). There were slight differences in the patterns among the stages within each treatment (Fig. 2). Additionally, cumulative survival did not differ among treatments at any stage (ANOVA, p > 0.162 in all cases).

**Figure 2:**
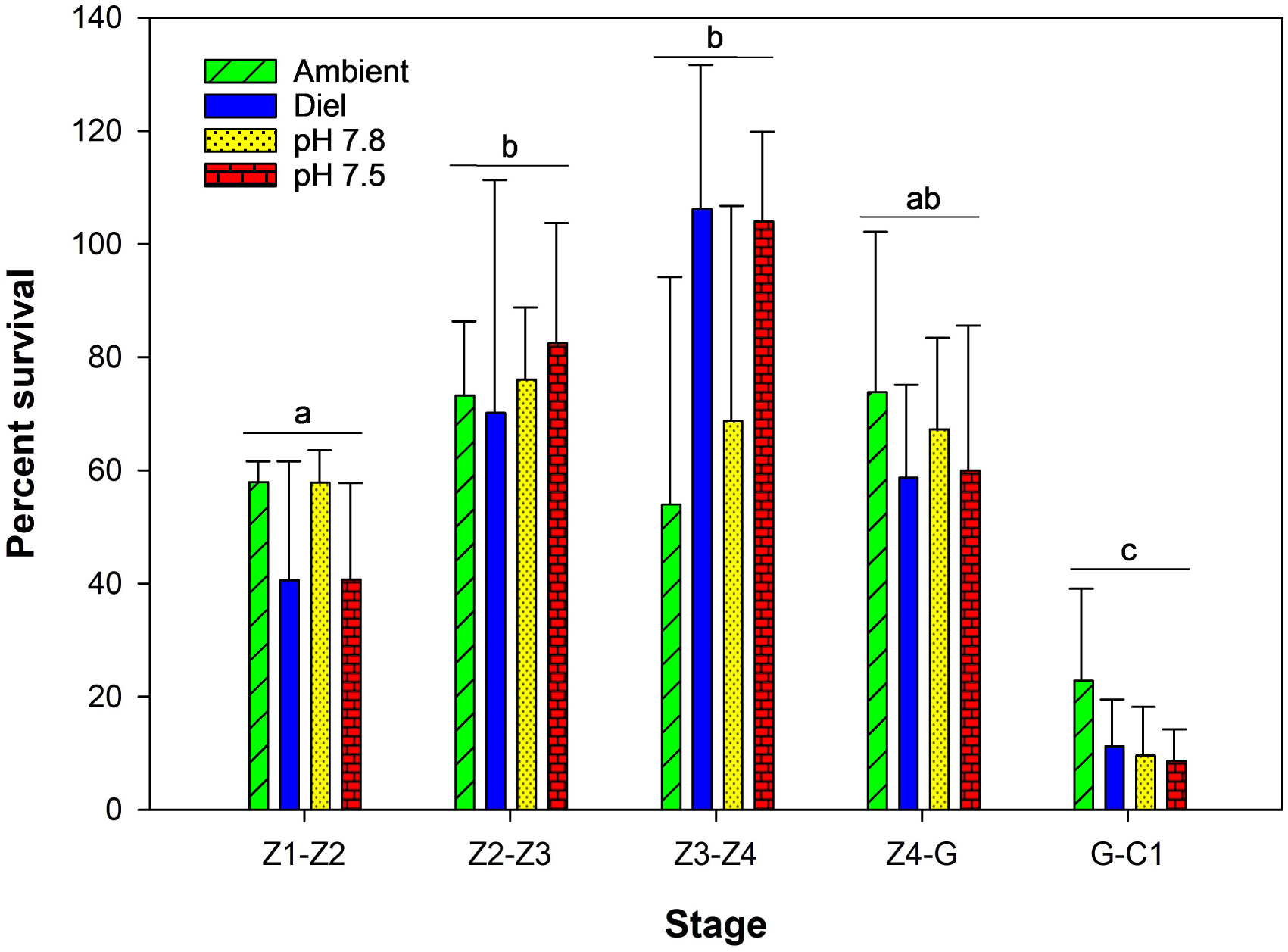
Survival of red king crab larvae between subsequent stages in four different pH treatments. Bars with different letters above them are significantly different (interactive effects are not shown here). Bars are means plus one standard deviation.

Dry mass increased with stage and there were differences among treatments but not their interaction (Table 2, Fig. 3). Overall, larvae in the Ambient treatment were slightly larger (effect size 7.8%) than larvae in pH 7.8 (Tukey’s test), but not pH 7.5 or Diel larvae; however, this pattern was reversed at the last stage where pH 7.8 crabs were 8% larger than the ambient (Fig. 3). C content differed among stages, treatments, and their interactions (Table 2, Fig. 4). In general, C content increased from the Z2 through the Z4 stage and then decreased slightly at the G stage and then by about 40% at the C1 stage (Fig. 3). There were no differences among pH treatments within any stage except that at the Z4 stage the Ambient larvae had a slightly lower C content then the other three. N content also differed among stages, treatments, and their interactions (Table 2, Fig. 4). The N content was relatively constant from the Z2-Z4 stages and then, similar to the C content dropping slightly at the G stage and substantially at the C1 stage (Fig. 4). There were no differences among pH treatments within any stage except that at the Z4 stage the Ambient larvae had a slightly lower N content then pH 7.8 and 7.5 treatments (Fig.4).

**Figure 3:**
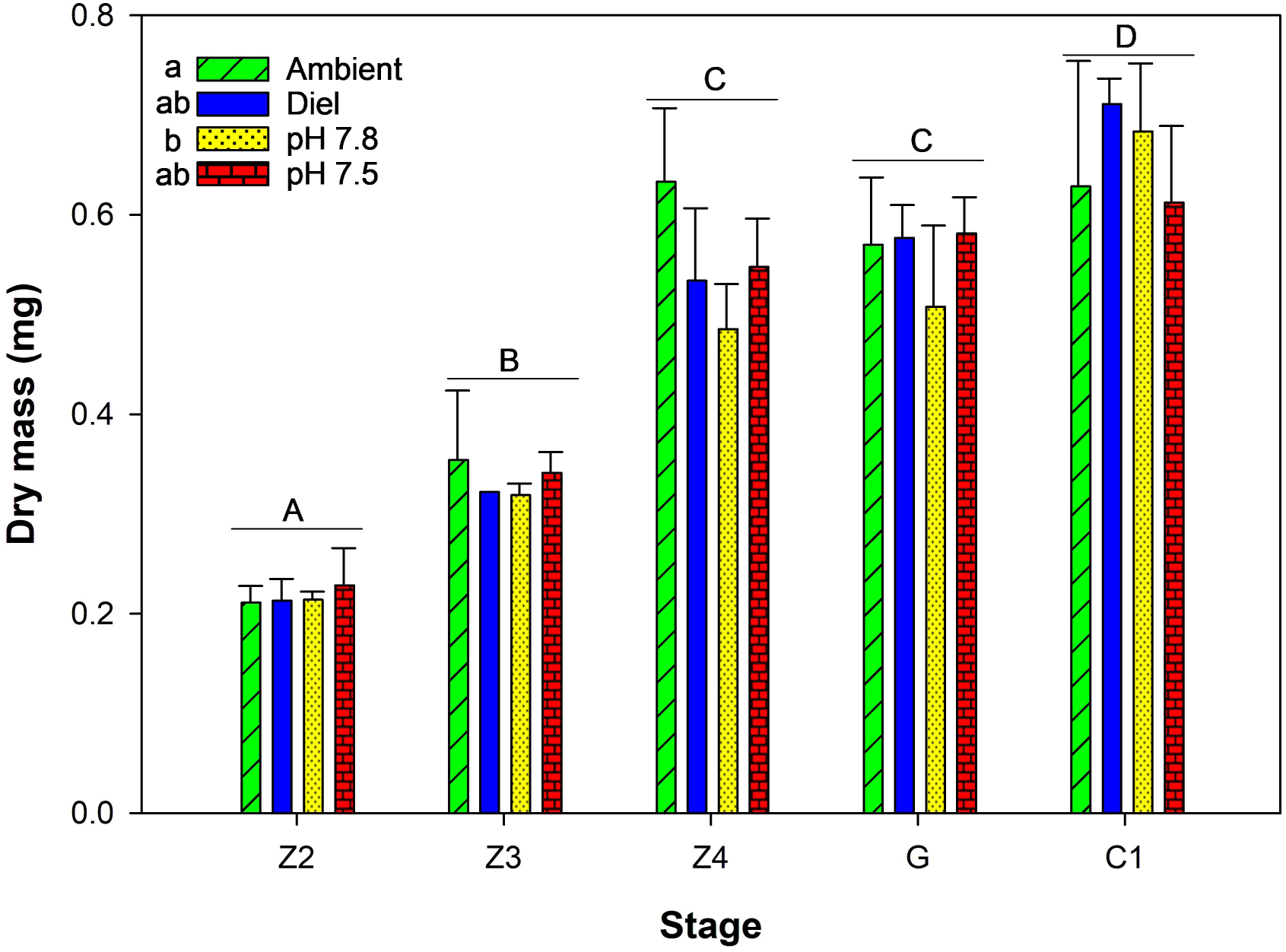
Dry mass of red king crab larvae between at each developmental stage in four different pH treatments. Bars or treatment indicated with different letters significantly different. Bars are means plus one standard deviation.

**Figure 4:**
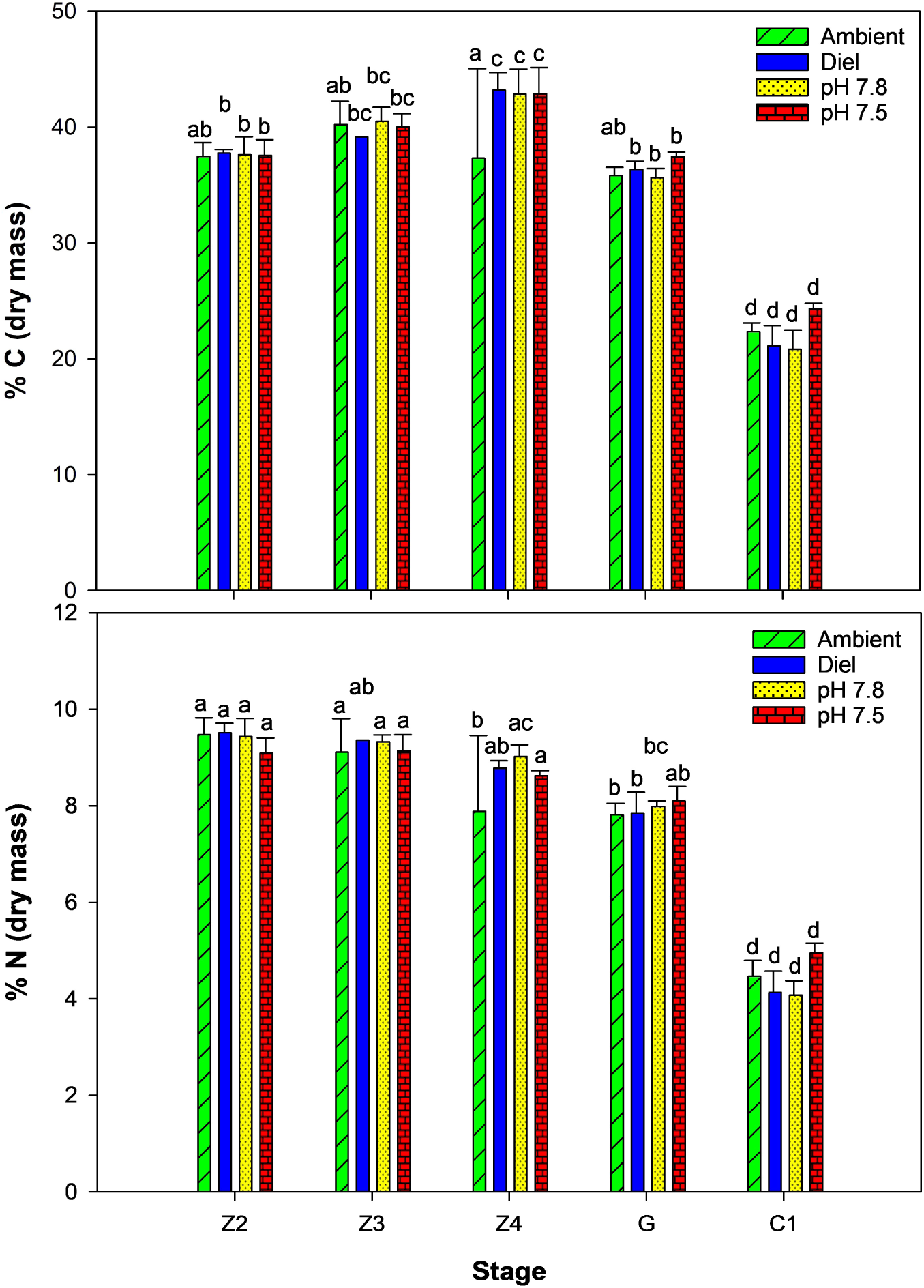
Carbon and nitrogen content of red king crab larvae at each developmental stage in four different pH treatments. Bars indicated with different letters significantly different. Bars are means plus one standard deviation.

Calcium content was low and did not differ among the zoeal stages, increased slightly at the G stage and greatly at the C1 stage and there was no difference among pH treatments (Table 2, Fig. 5). Magnesium content was lowest at the Z4 stage and highest and the C1 stage (Table 2, Fig. 5). Larvae in the ambient treatment had 20% higher magnesium content than the pH 7.8 and 7.5 treatments (Table 2, Fig. 5).

**Figure 5:**
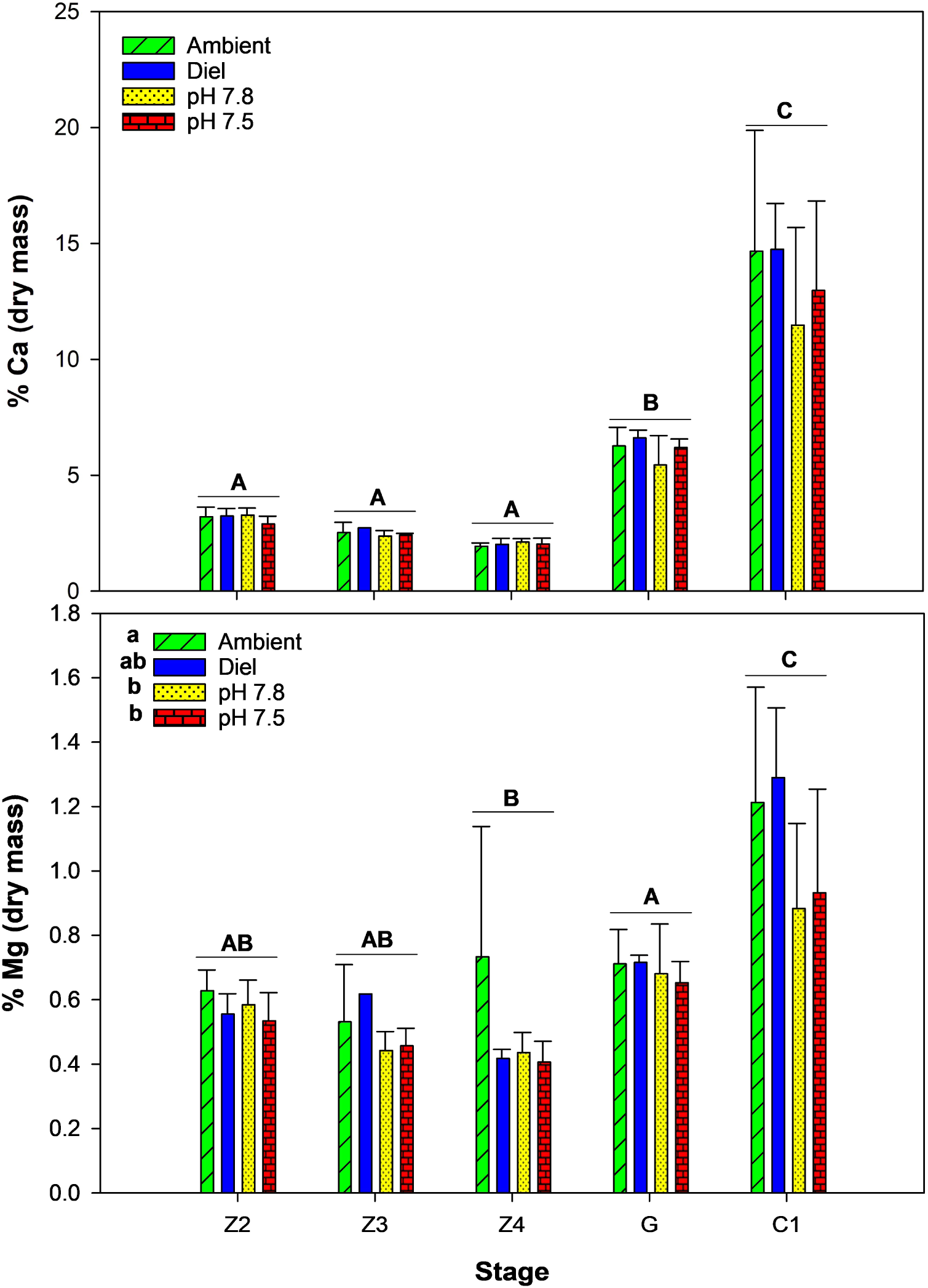
Calcium and magnesium content of red king crab larvae at each developmental stage in four different pH treatments. Bars indicated with different letters significantly different. Bars are means plus one standard deviation.

Morphometrics of the zoea different strongly among stages (PERMAOVA, Pseudo-F = 76.468, p < 0.00005) but not among pH treatments (Pseudo-F = 0.834, p = 0.530; Fig. 6). There was a significant interactive effect (Pseudo-F = 1.744, p = 0.034); however, in post-hoc comparisons there were no significant differences among treatments within each stage (p > 0.087 in all cases). Morphometrics of the G and C1 stages differed between stages (Pseudo-F = 76.468, p < 0.00005) and among treatments (Pseudo-F = 3.642, p = 0.012; Fig. 6). Post-hoc analyses indicated that only the ambient and pH 7.8 treatments differed from each other. We performed a similarity percentage (SIMPER) analysis to examine which features contributed to the differences; pH 7.8 crabs were smaller than ambient crabs in every measured variable with each contributing rough equally (16.2-25.0% range) to the distance between the groups. Effect sizes were small ranging from 0.9 to 8.0% (average 4.3%) among the measured variables.

**Figure 6:**
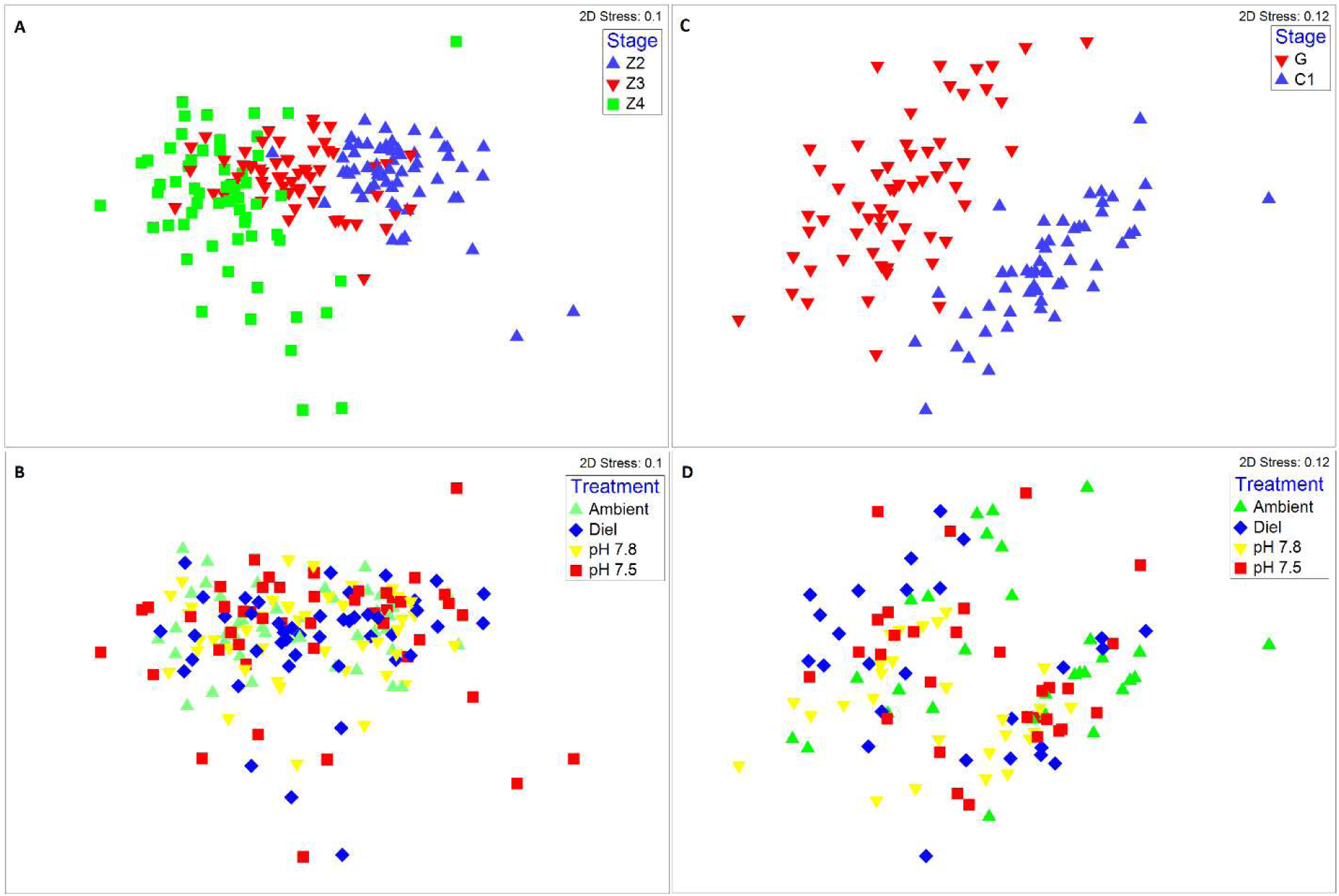
nMDS plots of red king crab zoeal (A & B) and glaucothoe and first crab stage (C & D) morphometrics from crabs reared in four pH treatments.

## Discussion

Red king crab larvae were highly resilient to changes in pH. In this study, larvae reared from hatching through settlement showed almost no effects of decreased pH. Most critically, there were no difference among treatments in either development time or survival. Where differences were detected they were almost always between the Ambient and pH 7.8 treatments, but, except for Mg content, there were no differences between Ambient and pH 7.5. Additionally, for the variables that did differ among treatments, effect sizes tended to be less than 5% or transitory, affecting one stage, but not others, and the reduced pH crabs were not always negatively affected. There were no differences between the Diel and Ambient treatments, suggesting that migration below the mixed layer under current conditions would not affect larval fitness. Taken together, this suggests that red king crab larvae are unlikely to be affected by ocean acidification for the foreseeable future.

Mortality varied among larval stages but not among pH treatments. In this study, survival rates increased from the Z1 to the Z3 stage, dropped for the Z4 stage and were quite low for the C1 stage. The ontogenetic pattern in mortality rates seems to differ greatly from study to study. At times there is a pattern of decreasing mortality throughout rearing with Z1s having only a 35% survival rate increasing to close to 100% from the glaucothoe to C1 molt (Sato and Tanaka, 1949b; Kovatcheva et al., 2006). In other studies mortality is lowest at early stages and lowest at the G and C1 stages (Nakanishi and Naryu, 1981; Nakanishi, 1985; Persselin and Daly, 2010; Swingle et al., 2013). Although these general trends were evident within individual studies, similar to our study, variance in stage-specific survival rates in larval rearing is high within each of the cited studies. In this study, survival rates to the C1 stage averaged 1.9% and ranged from 0 (in one replicate) to 4.5%. This is within the range of survival of larger scale hatchery-reared red king crab of a similar scale to our design, albeit on the low end (Nakanishi, 1976; Nakanishi and Naryu, 1981; Persselin and Daly, 2010; Swingle et al., 2013).

Similarly, development rate, growth rate, mass, and carbon and nitrogen content differed ontogenetically, but not with treatment pH. The developmental rate was on the slower side, but comparable to other studies (Sato and Tanaka, 1949b; Kurata, 1960; Kovatcheva et al., 2006; Long, 2016). Mass and linear size increased with stage as expected (Sato and Tanaka, 1949a; Nakanishi, 1987). C and N contents remained relatively constant throughout development, increasing slightly between the Z1 and Z4 stages, before sharply dropping between the G and C1 stage. This is almost certainly because glaucothoe are non-feeding and so during this stage the larvae were depleting their energy reserves.

However, there was no indication that pH substantially altered any of these metrics. Where differences occurred, they were typically between the Ambient and the pH 7.8, but significantly not the pH 7.5, and effect sizes were small and inconsistent among the stages, leading us to suspect type I statistical error.

Contrary to our expectations, there was no effect of water pH on survival either when survival at each stage was considered separately or cumulatively to any stage. Nor did pH affect the growth, size, shape, or carbon or nitrogen content of the larvae. In previous work on larvae (Long et al., 2013a) and juvenile red king crab (Long et al., 2013b; Swiney et al., 2017) both stages consistently showed higher mortality rate under low pH conditions. Starvation-survival times of red king crab larvae under acidified conditions were much lower than under ambient conditions and carbon content dropped faster suggesting that the larvae increased energetic demand to maintain acid-base homeostasis in low pH and depleted their energetic reserves faster (Long et al., 2013a). Similarly, red king crab juveniles increase their respiration rate under low pH conditions (Long et al., 2019) resulting in lower growth and higher mortality (Long et al., 2013b). Given these previous results, we expected to see depletion of energy reserves throughout larval development, slower growth, and higher mortality but did not. A likely explanation is that if energetic demand was higher, that the larvae were able to increase their feeding rate to compensate and thus maintain both homeostasis and growth rates. Such compensatory feeding is used by other species to reduce the effects of OA. For example, increasing food availability for larvae of the Olympia oyster, *Ostrea lurida*, reduces, but does not eliminate, the negative effects of OA on growth and successful metamorphosis (Hettinger et al., 2013). Similarly, larval copepods (*Tigriopus japonicus*) suffer greater reductions in development timing under reduced pH at lower food concentrations (Li et al., 2018) and juvenile corals (*Balanophyllia elegans*) are able to maintain calcification rates under reduced pH under food replete conditions (Crook et al., 2013). Future experiments examining feeding and respiration rates of red king crab larvae under reduced pH and experiments crossing food availability with pH would help to determine if this was indeed the case. Finally, although some crustaceans become deformed under reduced pH, either through dissolution (Kurihara et al., 2008) or through developmental disruptions (Agnalt et al., 2013), this was not the case for red king crab.

Stage-specific differences in the response to reduced pH are not uncommon among decapod crustaceans. Larval European lobsters, *Homarus gammarus*, for example, show a slight increase in mortality during the first larval stage only with no effects on growth at any stage (Small et al., 2015). In contrast, juveniles suffered increased mortality, decreased growth, decreased metabolism, and carapace deformation under reduced pH conditions (Agnalt et al., 2013; Small et al., 2016) likely driven by the high cost of maintaining acid-base hemostasis and calcification rates (Small et al., 2020). Similarly, American lobster, *Homarus americanus*, larvae were unaffected even by extreme low pH conditions whereas juveniles suffered increased mortality and decreased growth (Noisette et al., 2021). Although there are few decapod species for which comparable data is available to contrast the relative sensitivity of the larval and juvenile stages, it is interesting that for the three we are aware of, European and American lobsters and red king crab, the early benthic stage juveniles are more sensitive than the larvae in all cases. At least for the American lobster, metabolic profiling shows large changes with pH at the larval but not at the juvenile stages (Noisette et al., 2021) which suggests that the larval stages have a more plastic response to pH than do juveniles which imparts greater tolerance. Whether that is the case for red king crab is not certain. At the juvenile stage there is evidence for changes in gene expression relating to carapace formation and calcification (Stillman et al., 2020) as well as energetics (Spencer et al., In review), suggesting a level of plasticity at that stage at least; however, comparable data is not available for the larva. Future work should examine the mechanisms which convey tolerance to low pH conditions at the larval phase to compare to that of the juveniles.

Mineral content varied both with stage and pH treatment. In general, calcium and magnesium content were low during zoeal stages, increased moderately at the glaucothoe stage, and dramatically at the first crab stage. This ontogenetic change is consistent with the habitat and larval behavior. Zoeal stages are strictly planktonic and spend all of their time in the water column where their food source is. Strongly calcified cuticles would decrease buoyancy and increase the energetic cost of swimming up. Glaucothoe are an intermediate, semi-demersal stage that will swim to the bottom to look for suitable habitat and if not found, they will swim up into the water column again to move to a new location. Thus, they require a higher density and the increased protection of a harder cuticle likely imparts some anti-predator protection. Juvenile are fully benthic and fully calcified cuticles provide the needed negative buoyancy as well as protection from predation. Magnisium was lower in both the pH 7.8 and 7.8 treatments than in the ambient and the effect size, 27%, was largest at the C1 stage; there was no significant difference in calcium content. This shift suggests that although the crabs are maintaining their calcification rates under reduced pH conditions that there are changes to minerology. Many crustaceans, including red king crab (Long et al., 2013a; Long et al., 2013b), can maintain (Algayer et al., 2023) or even increase (Ries et al., 2009; Coffey et al., 2017; Glandon et al., 2018) their calcification rates under reduced pH conditions, likely because buffering their hemolymph with carbonate can help facilitate the calcification process (Ries et al., 2009; Whiteley, 2011). However, reduced pH can induce changes in mineral form, particularly between calcite and amorphous calcium carbonate and this change in magnesium content suggests this is occurring in this case (Dickenson et al., 2021). A higher Mg:Ca ratio in calcite increases the hardness (Magdans and Gies, 2004) but also its susceptibility to dissolution (Morse et al., 2006; Andersson et al., 2008) suggesting a potential tradeoff between these two functions.

Overall, red king crab larvae appear well adapted to low pH conditions, able to maintain their growth and survival rates over a wide range of conditions. Given the susceptibility of juveniles to low pH, this work highlights the large variance in the response of organisms across different life-history stages that can occur and reinforces that tolerance at one stage does not imply tolerance at all. Given the interactive effect in juveniles between temperature and pH (Swiney et al., 2017) future work should examine if increased temperature affects the response to pH in larvae. In addition, as ocean acidification can affect behavior in marine organisms, including crab, the effect on larval behavior and particularly the settling behavior of glaucothoe, should be investigated. Finally, as carryover effect from one life-history stage can affect the sensitively of subsequent stages (Hettinger et al., 2012), potential carryover effects from larval exposure on juvenile sensitive should be explored.

## Acknowledgements

This project was funded by the National Oceanic and Atmospheric Administration (NOAA) Ocean Acidification Program. We thank the staff of the Kodiak Laboratory for help performing the experiments. The findings and conclusions in the paper are those of the authors and do not necessarily represent the views of the National Marine Fisheries Service, NOAA. Reference to trade names or commercial firms does not imply endorsement by the National Marine Fisheries Service, NOAA.

## Notes

### Competing Interest Statement

The authors have declared no competing interest.

## Reference

Agnalt, A. L., Grefsrud, E. S., Farestveit, E., Larsen, M., and Keulder, F. 2013. Deformities in larvae and juvenile European lobster (*Homarus gammarus*) exposed to lower pH at two different temperatures. Biogeosciences, 10: 7883–7895, doi:10.5194/bg-10-7883-2013.

Algayer, T., Mahmoud, A., Saksena, S., Long, W. C., Swiney, K. M., Foy, R. J., Steffel, B. V., et al. 2023. Adult snow crab, Chionoecetes opilio, display body-wide exoskeletal resistance to the effects of long-term ocean acidification. Marine Biology, 170: 63, doi:10.1007/s00227-023-04209-0.

Andersson, A. J., Mackenzie, F. T., and Bates, N. R. 2008. Life on the margin: implications of ocean acidification on Mg-calcite, high latitude and cold-water marine calcifiers. Marine Ecology Progress Series, 373: 265–273, doi:10.3354/meps07639.

Bednaršek, N., Ambrose, R., Calosi, P., Childers, R. K., Feely, R. A., Litvin, S. Y., Long, W. C., et al. 2021. Synthesis of thresholds of ocean acidification impacts on decapods. Frontiers in Marine Science, 8: 651102, doi:10.3389/fmars.2021.651102.

Coffey, W. D., Yarram, A., Matoke, B., Long, W. C., Swiney, K. M., Foy, R. J., and Dickenson, G. 2017. Ocean acidification leads to altered micromechanical properties of the mineralized cuticle of juvenile red and blue king crabs. Journal of Experimental Marine Biology and Ecology, 495: 1–12, doi:doi.org/10.1016/j.jembe.2017.05.011.

Crook, E. D., Cooper, H., Potts, D. C., Lambert, T., and Paytan, A. 2013. Impacts of food availability and pCO_2_ on planulation, juvenile survival, and calcification of the azooxanthellate scleractinian coral *Balanophyllia elegans*. Biogeosciences, 10: 7599–7608, doi:10.5194/bg-10-7599-2013.

Daly, B., Eckert, G. L., and White, T. D. 2013. Predation of hatchery-cultured juvenile red king crabs (*Paralithodes camtschaticus*) in the wild. Canadian Journal of Fisheries and Aquatic Sciences, 70: 358–366, doi:10.1139/cjfas-2012-0377.

Dew, C. B. 1990. Behavioral ecology of podding red king crab, *Paralithodes camtschatica*. Canadian Journal of Fisheries and Aquatic Sciences, 47: 1944–1958.

Dickenson, G. H., Bejerano, S., Salvador, T., Makdisi, C., Patel, S., Long, W. C., Swiney, K. M., et al. 2021. Ocean acidification alters exoskeleton properties in adult Tanner crabs, *Chionoecetes baridi*. Journal of Experimental Biology, 224: jeb232819, doi:10.1242/jeb.232819.

Dickson, A. G., Sabine, C. L., and Christian, J. R. 2007. Guide to best practices for ocean CO_2_ measurements. PICES Special Publication 3: 191 p.

DOE. 1994. Handbook of methods for the analysis of the various parameters of the carbon dioxide system in sea water. Version 2. ORNL/CDIAC-74: 197 p.

Glandon, H. L., Kilbourne, K. H., Schijf, J., and Miller, T. J. 2018. Counteractive effects of increased temperature and pCO_2_ on the thickness and chemistry of the carapace of juvenile blue crab, *Callinectes sapidus*, from the Patuxent River, Chesapeake Bay. Journal of Experimental Marine Biology and Ecology, 498: 39–45.

Gnaiger, E., and Bitterlich, G. 1984. Proximate biochemical composition and caloric content calculated from elemental CHN analysis: a stoichiometric concept. Oecologia, 62: 289–298.

Haynes, E. B. 1983. Distribution and abundance of larvae of king crab, *Paralithodes camtschatica*, and pandalid shrimp in the Kachemak Bay area, Alaska, 1972 and 1976. NOAA Technical Report,N. O. a. A. A. U.S. Department of Commerce, National Marine Fisheries Service NMFS SSRF-765: 64 p.

Hettinger, A., Sanford, E., Hill, T. M., Hosfelt, J. D., Russell, A. D., and Gaylord, B. 2013. The influence of food supply on the response of Olympia oyster larvae to ocean acidification. Biogeosciences, 10: 6629–6638, doi:10.5194/bg-10-6629-2013.

Hettinger, A., Sanford, E., Hill, T. M., Russell, A. D., Sato, K. N. S., Hoey, J., Forsch, M., et al. 2012. Persistent carry-over effects of planktonic exposure to ocean acidification in the Olympia oyster. Ecology, 93: 2758–2768.

Kovatcheva, N. P., Epelbaum, A., Kalinin, A., Borisov, R., and Lebedev, R. 2006. Early life history stages of the red king crab Paralithodes camtschaticus (Tilesius, 1815): biology and culture, VNIRO Publishing, Moscow.

Kurata, H. 1960. Studies on the larvae and postlarvae of *Paralithodes camtschatica* III. The influence of temperature and salinity on the survival and growth of the larvae. Bulletin of the Hokkaido Regional Fisheries Research Laboratory, 21: 9–14.

Kurihara, H., Matsui, M., Furukawa, H., Hayashi, M., and Ishimatsu, A. 2008. Long-term effects of predicted future seawater CO_2_ conditions on the survival and growth of the marine shrimp *Palaemon pacificus*. Journal of Experimental Marine Biology and Ecology, 367: 41–46, doi:10.1016/j.jembe.2008.08.016.

Lavigne, H., and Gattuse, J. 2012. seacarb: Seawater carbonate chemistry with R. http://CRAN.R-project.org/package=seacarb. R package version 2.4.6 edn.

Li, F., Shi, J. H., Cheung, S. G., Shin, P. K. S., Liu, X. S., Sun, Y., and Mu, F. H. 2018. The combined effects of elevated pCO_2_ and food availability on *Tigriopus japonicus* Mori larval development, reproduction, and superoxide dismutase activity. Marine Pollution Bulletin, 126: 623–628, doi:10.1016/j.marpolbul.2017.02.054.

Long, W. C. 2016. A new quantitative model of multiple transitions between discrete stages, applied to the development of crustacean larvae. Fishery Bulletin, 114: 58–66, doi:10.7755/FB.114.1.5.

Long, W. C., Popp, J., Swiney, K. M., and Van Sant, S. B. 2012. Cannibalism in red king crab, *Paralithodes camtschaticus* (Tilesius, 1815): Effects of habitat type and predator density on predator functional response. Journal of Experimental Marine Biology and Ecology, 422-423: 101–106, doi:10.1016/j.jembe.2012.04.019.

Long, W. C., Pruisner, P., Swiney, K. M., and Foy, R. 2019. Effects of ocean acidification on respiration, feeding, and growth of juvenile red and blue king crabs (*Paralithodes camtschaticus* and *P. platypus*). ICES Journal of Marine Science, 76: 1335–1343, doi:10.1093/icesjms/fsz090.

Long, W. C., Swiney, K. M., and Foy, R. J. 2013a. Effects of ocean acidification on the embryos and larvae of red king crab, *Paralithodes camtschaticus*. Marine Pollution Bulletin, 69: 38–47, doi:doi.org/10.1016/j.marpolbul.2013.01.011.

Long, W. C., Swiney, K. M., and Foy, R. J. 2016. Effects of high *p*CO_2_ on Tanner crab reproduction and early life history, Part II: carryover effects on larvae from oogenesis and embryogenesis are stronger than direct effects. ICES Journal of Marine Science, 73: 836–848, doi:10.1093/icesjms/fsv251.

Long, W. C., Swiney, K. M., Harris, C., Page, H. N., and Foy, R. J. 2013b. Effects of ocean acidification on juvenile red king crab (*Paralithodes camtschaticus*) and Tanner crab (*Chionoecetes bairdi*) growth, condition, calcification, and survival. PloS one, 8: e60959, doi:doi.org/10.1371/journal.pone.0060959.

Long, W. C., Van Sant, S. B., and Haaga, J. A. 2015. Habitat, predation, growth, and coexistence: Could interactions between juvenile red and blue king crabs limit blue king crab productivity? Journal of Experimental Marine Biology and Ecology, 464: 58–67, doi:10.1016/j.jembe.2014.12.011.

Long, W. C., Van Sant, S. B., Swiney, K. M., and Foy, R. 2017. Survival, growth, and morphology of blue king crabs: Effect of ocean acidification decreases with exposure time. ICES Journal of Marine Science, 74: 1033–1041, doi:doi.org/10.1093/icesjms/fsw197.

Long, W. C., and Whitefleet-Smith, L. 2013. Cannibalism in red king crab: Habitat, ontogeny, and the predator functional response. Journal of Experimental Marine Biology and Ecology, 449: 142–148, doi:10.1016/j.jembe.2013.09.004.

Lyons, C., Eckert, G., and Stoner, A. W. 2016. Influence of temperature and congener presence on blue (*Paralithodes platypus* Brandt, 1850) and red (Paralithodes camtschaticus Tilesius, 1815) king crabs habitat preference and fish predation. Journal of Crustacean Biology, 36: 12–22.

Magdans, U., and Gies, H. 2004. Single crystal structure analysis of sea urchin spine calcites: Systematic investigations of the Ca/Mg distribution as a function of habitat of the sea urchin and the sample location in the spine. European Journal of Mineralogy, 16: 261.

Mathis, J. T., Cross, J. N., and Bates, N. R. 2011. Coupling primary production and terrestrial runoff to ocean acidification and carbonate mineral suppression in the eastern Bering Sea. Journal of Geophysical Research-Oceans, 116: 24, doi:10.1029/2010jc006453.

McMurray, G., Vogel, A. H., Fishman, P. A., Armstrong, D. A., and Jewett, S. C. 1986. Distribution of larval and juvenile red king crab (*Paralithoides camtschatica*) in Bristol Bay. Outer Continental Shelf Environmental Assessment Program: Final Reports of Principal Investigators, 53: 267–477 pp.

Meseck, S. L., Alix, J. H., Swiney, K. M., Long, W. C., Wikfors, G. H., and Foy, R. J. 2016. Ocean acidification affects hemocyte physiology in the Tanner crab (*Chionoecetes bairdi*). PloS one, 11: e0148477, doi:doi.org/10.1371/journal.pone.0148477.

Millero, F. J. 1986. The pH of estuarine waters. Limnology and Oceanography: 839–847.

Morse, J. W., Andersson, A. J., and Mackenzie, F. T. 2006. Initial responses of carbonate-rich shelf sediments to rising atmospheric pCO_2_ and “ocean acidification”: Role of high Mg-calcites. Geochimica et Cosmochimica Acta, 70: 5814–5830.

Nakanishi, T. 1976. Rearing larvae and post-larvae of the king crab (Paralithodes camtschatica). In FAO Technical Conference on Aquaculture, p. 3. Food and Agriculture Organization of the United Nations, Kyoto, Japan.

Nakanishi, T. 1985. The effects of the environment on the survival rate, growth and respiration of eggs, larvae and post-larvae of king crab (*Paralithodes camtschatica*). In Proceedings of the International King Crab Symposium, 5th edn, pp. 167–185. Ed. by S. K. Davis, F. Gaffney, J. McCrary, A. J. Paul, and R. S. Otto. Alaska Sea Grant College Program, University of Alaska Fairbanks, Anchorage, AK.

Nakanishi, T. 1987. Rearing condition of eggs, larvae and post-larvae of king crab. Bulletin of the Japan Sea Regional Fisheries Research Laboratory, 37: 57–161.

Nakanishi, T., and Naryu, M. 1981. Some aspects of large-scale rearing of larvae and postlarvae of the king crab (*Paralithodes camtschatica*). Bulletin of the Japan Sea Regional Fisheries Research Laboratory, 32: 39–47.

Noisette, F., Calosi, P., Madeira, D., Chemel, M., Menu-Courey, K., Piedalue, S., Gurney-Smith, H., et al. 2021. Tolerant larvae and sensitive juveniles: Integrating metabolomics and whole-organism responses to define life-stage specific sensitivity to ocean acidification in the American lobster. Metabolites, 11: 584, doi:10.3390/metabo11090584.

Pane, E. F., and Barry, J. P. 2007. Extracellular acid-base regulation during short-term hypercapnia is effective in a shallow-water crab, but ineffective in a deep-sea crab. Marine Ecology Progress Series, 334: 1–9, doi:10.3354/meps334001.

Persselin, S., and Daly, B. 2010. Diet and water source effects on larval red king crab cultivation. In Biology and Management of Exploited Crab Populations under Climate Change, 25th edn, pp. 479–494. Ed. by G. H. Kruse, G. L. Eckert, R. J. Foy, R. N. Lipcius, B. Sainte-Marie, D. L. Stram, and D. Woodby. Alaska Sea Grant College Program, University of Alaska Fairbanks, Anchorage, AK.

Pirtle, J. L., Eckert, G. L., and Stoner, A. W. 2012. Habitat structure influences the survival and predator-prey interactions of early juvenile red king crab *Paralithodes camtschaticus*. Marine Ecology Progress Series, 465: 169–184, doi:10.3354/meps09883.

Powell, G. C., and Nickerson, R. B. 1965. Aggregations among juvenile king crabs (*Paralithodes camtschatica*, Tilesius) Kodiak, Alaska. Animal Behaviour, 13: 374–380.

Punt, A. E., Dalton, M. G., Daly, B., Jackson, T., Long, W. C., Stockhausen, W. T., Szuwalski, C., et al. 2022. A framework for assessing harvest strategy choice when considering multiple interacting fisheries and a changing environment: The example of eastern Bering Sea crab stocks. Fisheries Research, 252: 106338, 10.1016/j.fishres.2022.106338.

Punt, A. E., Poljak, D., Dalton, M. G., and Foy, R. J. 2014. Evaluating the impact of ocean acidification on fishery yields and profits: The example of red king crab in Bristol Bay. Ecological Modelling, 285: 39–53.

Ries, J. B., Cohen, A. L., and McCorkle, D. C. 2009. Marine calcifiers exhibit mixed responses to CO_2_-induced ocean acidification Geology, 37: 1131–1134.

Sato, S., and Tanaka, S. 1949a. Study on the larval stage of *Paralithodes camtschatica* (Tilesius) I. About morphological research. Bulletin of the Hokkaido Regional Fisheries Research Laboratory, 1: 7–24.

Sato, S., and Tanaka, S. 1949b. Study on the larval stage of *Paralithodes camtschatica* (Tilesius) II. On the rearing. Scientific papers of the Hokkaido Fisheries Science Institution, 3: 18–30.

Small, D., Calosi, P., White, D., Spicer, J. I., and Widdicombe, S. 2010. Impact of medium-term exposure to CO_2_ enriched seawater on the physiological functions of the velvet swimming crab *Necora puber*. Aquatic Biology, 10: 11–21, doi:10.3354/ab00266.

Small, D. P., Calosi, P., Boothroyd, D., Widdicombe, S., and Spicer, J. I. 2015. Stage-specific changes in physiological and life-history responses to elevated temperature and Pco_2_ during the larval development of the European lobster *Homarus gammarus* (L.). Physiological and Biochemical Zoology, 88: 494–507, doi:10.1086/682238.

Small, D. P., Calosi, P., Boothroyd, D., Widdicombe, S., and Spicer, J. I. 2016. The sensitivity of the early benthic juvenile stage of the European lobster *Homarus gammarus* (L.) to elevated pCO_2_ and temperature. Marine Biology, 163: 1–12.

Small, D. P., Calosi, P., Rastrick, S. P. S., Turner, L. M., Widdicombe, S., and Spicer, J. I. 2020. The effects of elevated temperature and *P*CO_2_ on the energetics and haemolymph pH homeostasis of juveniles of the European lobster, *Homarus gammarus*. The Journal of experimental biology, 223: jeb209221, doi:10.1242/jeb.209221.

Spencer, L., Long, W. C., Spies, I. B., Nichols, K., and Foy, R. In review. Narrowed gene functions and enhanced transposon activity are associated with high tolerance to ocean acidification in a subarctic crustacean.

Stevens, B. G. 2003. Settlement, substratum preference, and survival of red king crab *Paralithodes camtschaticus* (Tilesius, 1815) glaucothoe on natural substrata in the laboratory. Journal of Experimental Marine Biology and Ecology, 283: 63–78.

Stevens, B. G., and Kittaka, J. 1998. Postlarval settling behavior, substrate preference, and time to metamorphosis for red king crab *Paralithodes camtschaticus*. Marine Ecology Progress Series, 167: 197–206.

Stevens, B. G., and Swiney, K. M. 2005. Post-settlement effects of habitat type and predator size on cannibalism of glaucothoe and juveniles of red king crab *Paralithodes camtschaticus*. Journal of Experimental Marine Biology and Ecology, 321: 1–11, doi:10.1016/j.jembe.2004.12.026.

Stillman, J. H., Fay, S. A., Ahmad, S. M., Swiney, K. M., and Foy, R. J. 2020. Transcriptomic response to decreased pH in adult, larval and juvenile red king crab, *Paralithodes camtschaticus*, and interactive effects of pH and temperature on juveniles. Journal of the Marine Biological Association of the United Kingdom, 100: 251–265, doi:10.1017/s002531541900119x.

Swiney, K. M., and Long, W. C. 2015. Primiparous red king crab, *Paralithodes camtschaticus*, are less fecund than multiparous crab. Journal of Shellfish Research, 34: 493–498, doi:10.2983/035.034.0233.

Swiney, K. M., Long, W. C., Eckert, G. L., and Kruse, G. H. 2012. Red king crab, *Paralithodes camtschaticus*, size-fecundity relationship, and inter-annual and seasonal variability in fecundity. Journal of Shellfish Research, 31: 925–933.

Swiney, K. M., Long, W. C., and Foy, R. J. 2016. Effects of high *p*CO_2_ on Tanner crab reproduction and early life history-Part I: long-term exposure reduces hatching success and female calcification, and alters embryonic development ICES Journal of Marine Science, 73: 825–835, doi:doi.org/10.1093/icesjms/fsv201.

Swiney, K. M., Long, W. C., and Foy, R. J. 2017. Decreased pH and increased temperatures affect young-of-the-year red king crab (*Paralithodes camtschaticus*). ICES Journal of Marine Science, 74: 1191–1200, doi:10.1093/icesjms/fsw251.

Swingle, J. S., Daly, B., and Hetrick, J. 2013. Temperature effects on larval survival, larval period, and health of hatchery-reared red king crab, *Paralithodes camtschaticus*. Aquaculture, 384–387: 13–18, doi:10.1016/j.aquaculture.2012.12.015.

Wainwright, T. C., Armstrong, D. A., Andersen, H., Dinnel, P. A., Herren, D., Jensen, G. C., Orensanz, J. M., et al. 1991. Port Moller king crab studies. Annual Report, FRI-UW-9203: 38 p.

Walther, K., Anger, K., and Portner, H. O. 2010. Effects of ocean acidification and warming on the larval development of the spider crab *Hyas araneus* from different latitudes (54 degrees vs. 79 degrees N). Marine Ecology Progress Series, 417: 159–170, doi:10.3354/meps08807.

Whiteley, N. M. 2011. Physiological and ecological responses of crustaceans to ocean acidification. Marine Ecology Progress Series, 430: 257–271, doi:10.3354/meps09185.

